# Using CRISPR barcoding as a molecular clock to capture dynamic processes at single-cell resolution

**DOI:** 10.1101/2024.10.14.618192

**Authors:** Yolanda Andres-Lopez, Alice Santambrogio, Ioannis Kafetzopoulos, Christopher Todd, Carla El Khouri, Celia Alda-Catalinas, Stephen J. Clark, Wolf Reik, Irene Hernando-Herraez

**Author notes:** Joint Authors (Yolanda Andres-Lopez, Alice Santambrogio, Ioannis Kafetzopoulos).

## Abstract

Biological processes are fundamentally dynamic, yet existing methods for capturing these temporal changes are limited. We present scDynaBar, a novel approach that integrates CRISPR-Cas9 dynamic barcoding with single-cell sequencing to enable the recording of temporal cellular events. In this system, genetic barcodes accumulate mutations over a 25-day period and then are sequenced together with the transcriptome of each single cell. We propose that this gradual accumulation of genetic diversity can be exploited to create an ordered record of a cellular event. We apply this approach to track the transition from a pluripotent state to a two-cell (2C)–like state in mouse embryonic stem cells (mESCs). The results provide compelling evidence for the transient nature of the 2C-like state. Additionally, our system shows consistent mutation rates across diverse cell types in a mouse gastruloid model, underscoring its robustness and versatility across various biological contexts. This technique not only improves our ability to study single-cell dynamics but also creates new opportunities for recording other temporal signals—in other words, using dynamic barcoding as a molecular clock in individual cells.

**GRAPHICAL ABSTRACT:** 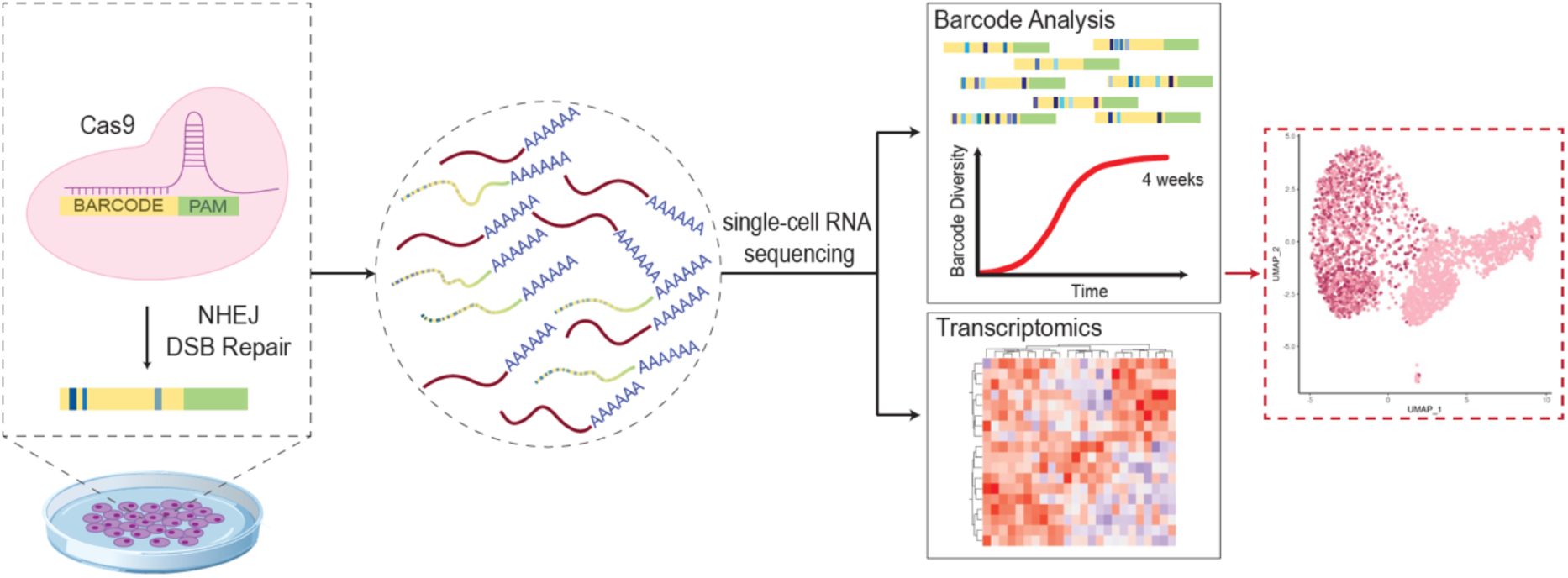

## INTRODUCTION

Cellular behaviour is intrinsically dynamic, undergoing alterations in response to internal and external stimuli. Different methodologies are commonly used to track or infer these changes. However, these methods present several limitations. For example, methods based on live-cell fluorescence microscopy are often incompatible with *in vivo* studies and are restricted in terms of long-term and throughput capabilities (*1*). Other methods, such as those based on inference algorithms using single-cell RNA sequencing (scRNA-seq) have low temporal resolution, as scRNA-seq only captures a snapshot of gene expression at one specific time point (*2*, *3*).

DNA memory systems based on CRISPR technology have emerged as valuable tools for recording dynamic events (*4–6*). These systems exploit CRISPR-based techniques to induce permanent genetic modifications at precise genomic locations. By temporally regulating the delivery of CRISPR components, these systems can generate DNA mutations that function as molecular barcodes, enabling retrospective analysis of cellular activities. A variety of DNA memory devices have been developed to track a wide range of biological signals (*4*), including gene expression (*7*, *8*), chemical exposure (*8–10*), lineage of origin (*11–13*), clonality (*14*, *15*), activation of signalling pathways (*16*) and inflammatory responses (*17*, *18*). Furthermore, several methods now enable single-cell resolution readouts. These approaches typically employ guide RNAs (gRNAs) targeting polyadenylation (polyA) sites, allowing simultaneous sequencing of both single-cell transcriptomes and DNA barcodes (*19*, *20*). This integrated strategy has been successfully applied to track lineage of origin in developmental (*21–27*) and cancer studies (*28–30*). More recently, prime editing has also been applied to HEK293T cells to track their lineage of origin at the single cell level (*15*).

Interestingly, dynamic CRISPR barcoding has also been proposed as a strategy for encoding time (*18*). This approach relies on self-targeting gRNAs, a variant of the canonical form of CRISPR-Cas9 gRNAs with the unique ability to repeatedly target their own sequence (*12*, *17*). The continuous mutagenesis leads to a gradual accumulation of mutations over time, which can be interpreted as a molecular record of elapsed time (*18*). However, since the target site is the gRNA locus itself that lacks polyadenylation, this dynamic barcoding approach is not compatible with standard single-cell RNA sequencing techniques.

In this study, we present scDynaBar, a novel method based on self-targeting gRNAs that enables 25-day barcoding with high recovery efficiency for both genetic barcodes and transcriptomes of individual cells using standard sequencing technologies. By coupling barcoding with the expression of Zscan4c, we successfully demonstrate the versatility of scDynaBar in tracking the transition from 2C-like states to pluripotency in mESCs *in vitro*. This approach revealed that the 2C-like state is transient and does not persist over extended periods. Additionally, our research shows consistent mutation rates in mouse gastruloids, suggesting similar DNA repair rates among different cell types. These findings highlight the broad potential of scDynaBar for tracking biological processes at single-cell resolution.

## RESULTS

### Dynamic barcoding

We aimed to develop a versatile system that can (i) record molecular events over extended periods, and (ii) simultaneously profile genetic barcodes and gene expression at the single-cell level, while being compatible with standard commercial sequencing technologies for easy implementation. To achieve this, we engineered self-targeting gRNAs (*17*) so that they can be retrieved from a polyadenylated transcript (expressed barcode). We also included an mCherry reporter which enables easy visualization and selection of targeted cells (**Figure 1A**).

**Figure 1.**
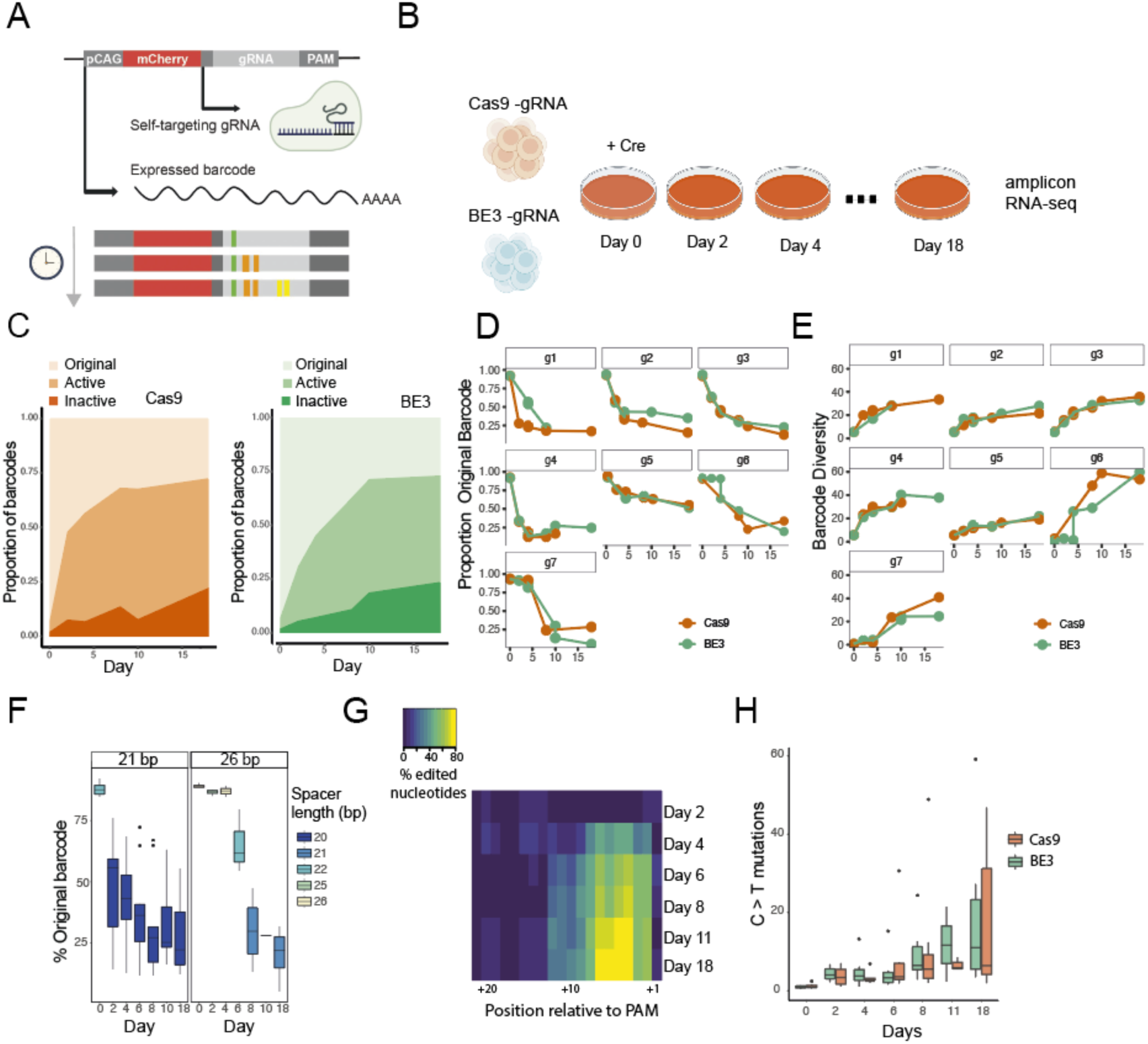
**A.** Schematic representation of dynamic barcoding with a poly-A readout. **B.** Schematic representation of the experimental setup. **C.** Proportion of barcodes based on their mutational profiles for Cas9 (left) and BE3 (right). Barcodes are grouped into three categories: edits on the protospacer with intact PAM motif (active); absence of PAM motif (inactive); and uncut gRNA (original). **D.** Proportion of original barcodes over time across different gRNAs. **E.** Barcode diversity over time, considering mismatches, gaps and gap extensions across different gRNAs. **F.** Proportion of original barcodes over time for gRNAs with 21-bp spacers (left) or 26-bp spacers (right). Box plots are coloured by the mean spacer length at different time points (Cas9 system). **G.** Percentage of the original nucleotide over time, aligning the spacer relative to the PAM sequence (Cas9 system). **H.** Percentage of C>T mutations over time, considering all different gRNAs classified by Cas9 versions. For all boxplots, the boxes represent the interquartile range (IQR), with the horizontal line inside each box indicating the median.

To investigate the dynamics of the system, and to test its ability to detect the barcodes at the RNA level, we performed a time-course experiment using seven different gRNAs with varying protospacer sizes (21 or 26 base pairs [bp]) and nucleotide compositions (**Supplementary Figure 1**). We also aimed to evaluate the relative efficacies of barcoding using standard Cas9 and a base editing system, as the latter avoids inducing double-strand breaks. Specifically, we used the BE3 base editor, which consists of a Cas9 nickase fused to the cytidine deaminase APOBEC and the uracil DNA glycosylase inhibitor (UGI) domain (*32*). The UGI domain facilitates uracil retention, ensuring precise C:G to T:A substitutions within a 4 to 8 nucleotide window (*32*). In its absence, the frequency of indels increases, along with a higher incidence of other substitutions (*32*, *33*). Therefore, to enhance mutational diversity in our approach, we used a BE3 system lacking this domain (APOBEC–XTEN–dCas9(A840H)) (*32*). Both Cas9 constructs contained a constitutive promoter (CMV early enhancer fused to modified chicken β-actin promoter) and a GFP reporter. Expression of the constructs was controlled by a Cre-responsive FLEX switch, ensuring that expression occurred only after recombination. We co-transfected individual gRNAs together with one of the two Cas9 constructs into mESCs. After stable integration, we activated Cas9 by adding Cre recombinase to the culture media and then collected cells from this mixed polyclonal population at various time points, up to day 18 (**Figure 1B**). We then selected cells positive for both reporters (GFP and mCherry) by fluorescence-activated cell sorting (FACS) and subsequently performed bulk amplicon sequencing on their RNA (see **Supplementary Figure 2 and Methods**) (**Figure 1B**).

To investigate the general dynamics of the systems, we first classified the barcode sequences into three groups: (1) the original barcode, in which no editing occurred, (2) an active barcode, in which editing occurred in the protospacer region but with the PAM sequence still uncut, and (3) an inactive barcode, in which editing occurred in both the protospacer regions and the PAM sequence, preventing further mutagenesis. Our data are consistent with sequential editing, as they show increased level of active and inactive gRNAs, and a decreased frequency of original barcodes, over time (**Figure 1C**). The increased editing rate is consistent when using any of the seven gRNAs that we tested, as well as with either version of Cas9 (**Figure 1D**) (Jonckheere-Terpstra test; p = 2 × 10^−13^). Background editing in the absence of Cre (at day 0) was low (averaging 3% across experiments), likely resulting from spontaneous Cas9 expression (**Supplementary Figure 3**). After 18 days of editing, the system still contained barcodes with no mutations (mean original barcode frequency, 26.8%), indicating that mutagenesis could potentially be extended beyond this day (**Figure 1D**). We next assessed the diversity of the edited reads by comparing their sequences to the original barcode. In this analysis, we performed pairwise alignments for over 100,000 barcode sequences and calculated a diversity score considering mismatches, gaps and gap extensions. Similarly, all seven gRNAs demonstrated a sequential increase in diversity over the 18 days (Jonckheere-Terpstra test; *p* = 8 × 10^−16^), providing strong evidence for continuous mutagenesis as well as for their capacity to effectively record molecular events (**Figure 1E, Supplementary Figure 4**). In agreement with previous studies, we also observed that in the Cas9 system 26-bp gRNAs exhibit a slower editing rate compared to 21 -bp gRNA during the initial four days (**Figure 1F**) (*31*) (Day 4, Mann-Whitney-Wilcoxon test; p = 0.0005). After this time point, the frequency of intact barcodes for both the 21-bp and the 26-bp barcodes became very similar. This could be attributed to the accumulation of deletions in the 26-bp gRNAs (**Figure 1F**), potentially leading to an increased editing rate at later time points.

Next, we analysed the distribution of mutagenesis relative to the PAM of each gRNA in the Cas9 system. As expected from the activity of Cas9, we observed a preferential accumulation of mutations near the cut site, specifically within three nucleotides upstream of the PAM site (**Figure 1G**). We then compared the specific types of edits in the BE3 system to the standard Cas9. Notably, we observed a similar accumulation of indels over time in both systems (Mann-Whitney-Wilcoxon test; p = 0.45, **Supplementary Figure 5**). This could be attributed to the absence of the UGI domain, which may lead to DNA repair by alternative cellular mechanisms, ultimately resulting in indel formation. Despite the lack of the UGI domain, our BE3 system exhibited a slightly higher frequency of C>T mutations at most time points, with an average difference of 6.9% (**Figure 1H**).

### Reproducibility and window of activity

We next aimed to extend the duration of mutagenesis to determine the maximum period during which the system can remain active and continue to generate mutations. As before, each of the seven gRNAs was individually co-transfected with one of the Cas9 constructs and then processed at various time points, up to day 32 (**Figure 2A**). We then compared original barcode frequencies across samples with two biological replicates (considering gRNA, day, and experiment, n=86). Remarkably, we observed consistent results between biological replicates for the initial 18 days, with a Pearson’s correlation coefficient of 0.94 (*p* < 2.2 × 10^−16^), indicating the high reproducibility of our system (**Figure 2B**). These results also indicate that the Cas9 system remained active until day 25, albeit working at a slower rate (**Figure 2C, D**). In contrast, the BE3 system exhibited a significant increase in the frequency of original barcodes and a decrease in sequence diversity at day 25 (**Figure 2C, D**). We hypothesize that this may result from differences in cellular fitness. Cells with high editing levels, particularly those with mutations in the PAM sequence, may no longer target the gRNA locus, leading to potential APOBEC-mediated off-target effects that could cause selective depletion (*34*). In contrast, cells with an intact PAM sequence would continue to recruit APOBEC to the gRNA locus, potentially reducing off-target effects and promoting their enrichment over time. Nonetheless, future experiments are needed to better understand this phenomenon.

**Figure 2.**
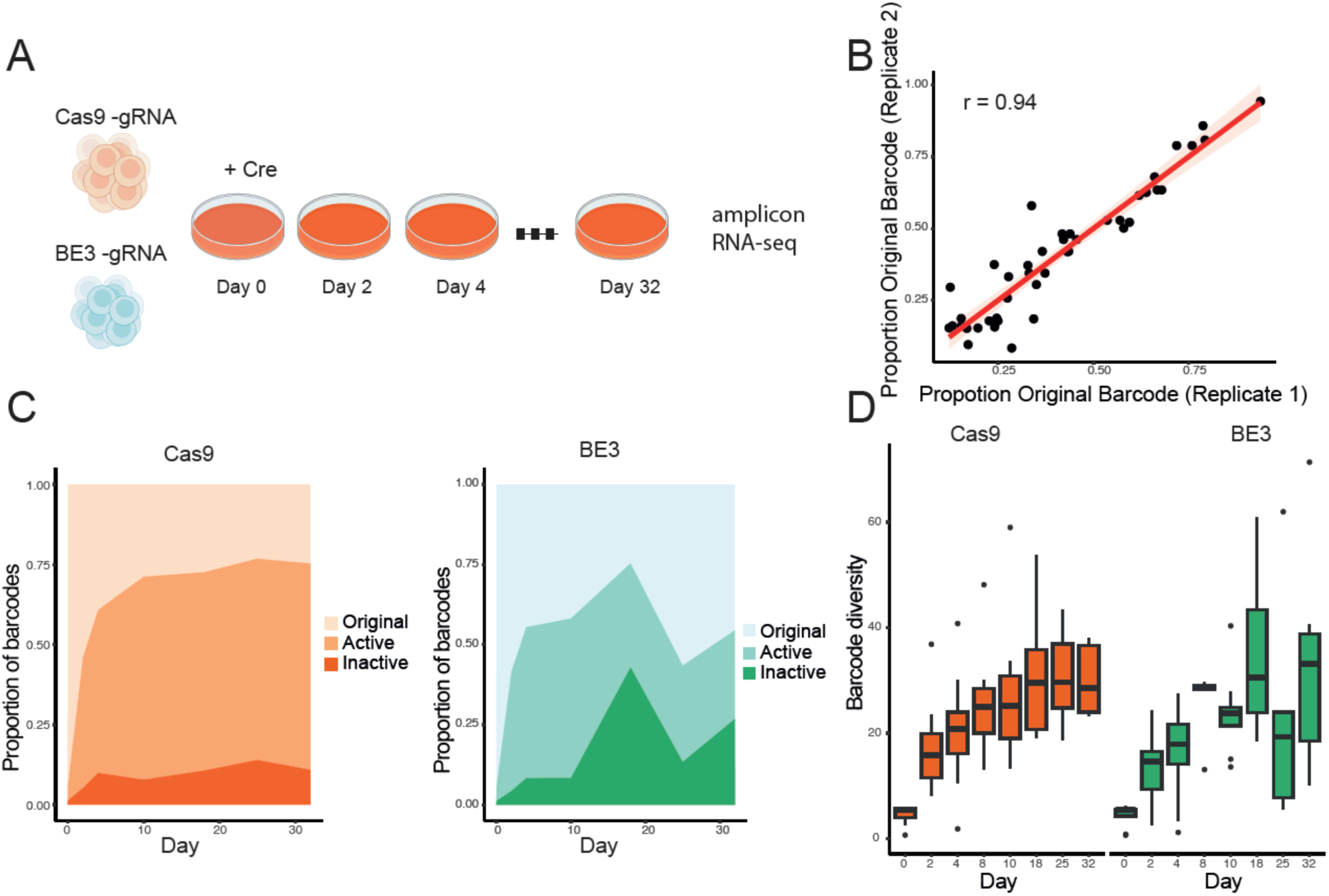
**A**. Schematic representation of the experimental setup. **B**. Pearson’s correlation of original barcode frequencies between biological replicates from days 0 to 18. Each point represents the correlation coefficient between biological replicates considering gRNA, experiment type and time point. A total of 86 samples were analysed; including 50 samples from theCas9 experiment and 36 from the BE3 experiment. **C**. Proportion of barcodes based on their mutational profile: edits on the protospacer with intact PAM motif (active); lack of PAM motif (inactive); and uncut gRNA (original) according to Cas9 (left) or BE3 (right). **D.** Barcode diversity over time, considering mismatches, gaps and gap extensions, pooling all gRNA for Cas9 (left) and BE3 (right). In the boxplots, the boxes represent the IQR with the horizontal line inside each box indicating the median.

In conclusion, we demonstrate that both systems are effective for sequential barcoding, with BE3 achieving comparable levels of barcode diversity in experiments conducted over a two-week period and Cas9 being more suitable for long-term applications.

### Simultaneous detection of genetic barcodes in single cells

We then evaluated whether genetic barcodes could be simultaneously recovered together with transcriptomic data using standard single-cell RNA sequencing. First, based on our previous observations, we selected gRNA g3 in combination with the canonical form of Cas9. We then established a clonal cell line to minimize potential variability due to different integration sites between individual cells. Using qPCR on genomic DNA, we determined that this clonal cell line contains a single integration (**Methods**, **Supplementary Figure 6).** We grew these cells under serum/LIF conditions and following Cre induction, we FACS-sorted double-positive cells (for GFP and mCherry) on days 4 and 10 (**Figure 3A**). Additionally, we included a non-induced group that showed minimal Cas9-GFP expression at day 10 (GFP = 0.1%), which confirmed low levels of background activity.

**Figure 3.**
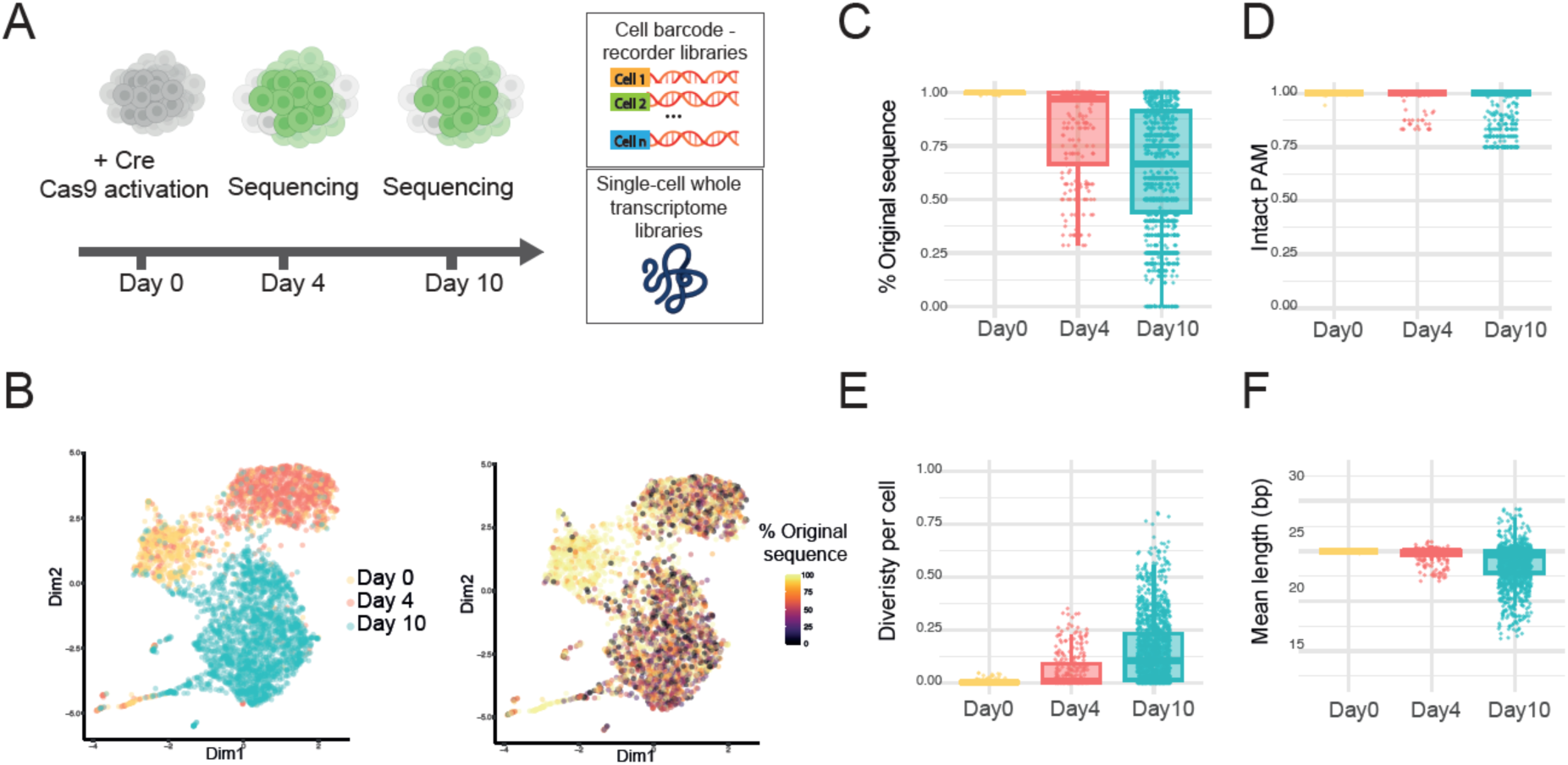
**A.** Schematic representation of the experimental setup. **B**. t-SNE plot coloured by cluster: non-induced (day 0) and induced (day 4 or day 10) (left). Colour indicates the mean percentage of the original barcode sequence per cell (right). **C–F**. Boxplot of results at days 0, 4 and 10 showing **C.** the percentage of the original barcode sequences per cell; **D**. the mean proportion of barcodes with full PAM sequence per cell; **E**. mean sequence diversity per cell and **F**. mean length of the barcodes per cell. For all boxplots, the boxes represent the IQR, with the horizontal line inside each box indicating the median. Each dot represents an individual cell, with data points displayed within the 95% confidence interval.

Cells were processed using the 3’ 10X Genomics platform, where they were encapsulated in droplets with barcoded PolyT-coated beads and subjected to whole transcriptome amplification following the standard 10x Genomics scRNA-seq 3′ protocol. We then specifically amplified and sequenced the genetic barcodes from the cell-barcoded cDNA (**Figure 3A**). Overall, we successfully recovered genetic barcodes from 97% of cells that passed transcriptome quality control, demonstrating the efficiency of our approach for simultaneous transcriptome and barcode detection. t-SNE analysis revealed three clusters of cells, corresponding to the different time points: non-induced (day 0; *n* = 495), induced at day 4 (*n* = 626) and induced at day 10 (*n* = 1117) (**Figure 3B**). The most frequent barcodes sequences are listed in **Supplementary Table 1**, illustrating the progressive increase in edited barcodes over time. For each cell, we assessed the level of barcoding by analysing: the proportion of reads carrying the original barcode sequence, the mean barcode diversity score (calculated as described previously), the average length of the barcode sequence, and the percentage of barcodes carrying a PAM sequence (**Figure 3C–F**). Our findings align with the results in bulk, showing sequential editing over time (Jonckheere-Terpstra test; *p* = 2 × 10^−04^, 1000 permutations). We observed a decrease in the presence of original sequences along with a decline of the proportion of barcodes carrying a PAM motif (**Figure 3C, D**). Additionally, we observed an increase in barcode diversity and greater variability in their lengths (**Figure 3E, F),** along with increased number of unique barcode sequences per cell **(Supplementary Figure 7).** We also observed that the 1,480 edited gRNAs sequences were unlikely to have off-targets effects that could affect cell viability (**Supplementary Table 2**). Altogether, these data demonstrate the effectiveness of dynamic barcoding in single cells and the capability of the method to simultaneously capture barcodes and transcriptomic information using standard single-cell technologies.

### Recording dynamic transitions

Having established the efficacy of scDynaBar in single cells, we next applied it to track transitions between pluripotency and totipotency-like states. Under standard serum/LIF culture conditions, a small population (less than 1%) of mESCs spontaneously emerges that exhibits features resembling the totipotent 2C stage (*35*, *36*). These features include a specific transcriptional profile, such as the expression of Zscan4 genes, increased histone mobility, DNA demethylation and the capacity to contribute to both embryonic and extraembryonic tissues (*35*, *37*, *38*). Although it is believed that the 2C-like state cannot self-propagate but rather continuously undergoes transitions to a pluripotent state, these dynamics have not been directly measured (*35*, *39*). To track 2C-like conversions, we integrated our barcoding approach with a pZscan4-CreERT2 construct (**Figure 4A**) (*40*). With this setup, cells entering the 2C-like state permanently activate the barcoding cassette during treatment with tamoxifen. After establishing a monoclonal line, cells were treated with tamoxifen for 12 days. Following treatment, we observed an increase in GFP-positive cells (9% tamoxifen-treated cells; 0.1% in untreated cells). Double-positive cells (GFP- and mCherry-positive) were FACS-sorted and processed for single-cell RNA-seq and barcoding sequencing (**Figure 4A**). We also sequenced a portion of GFP-negative cells as a control.

**Figure 4.**
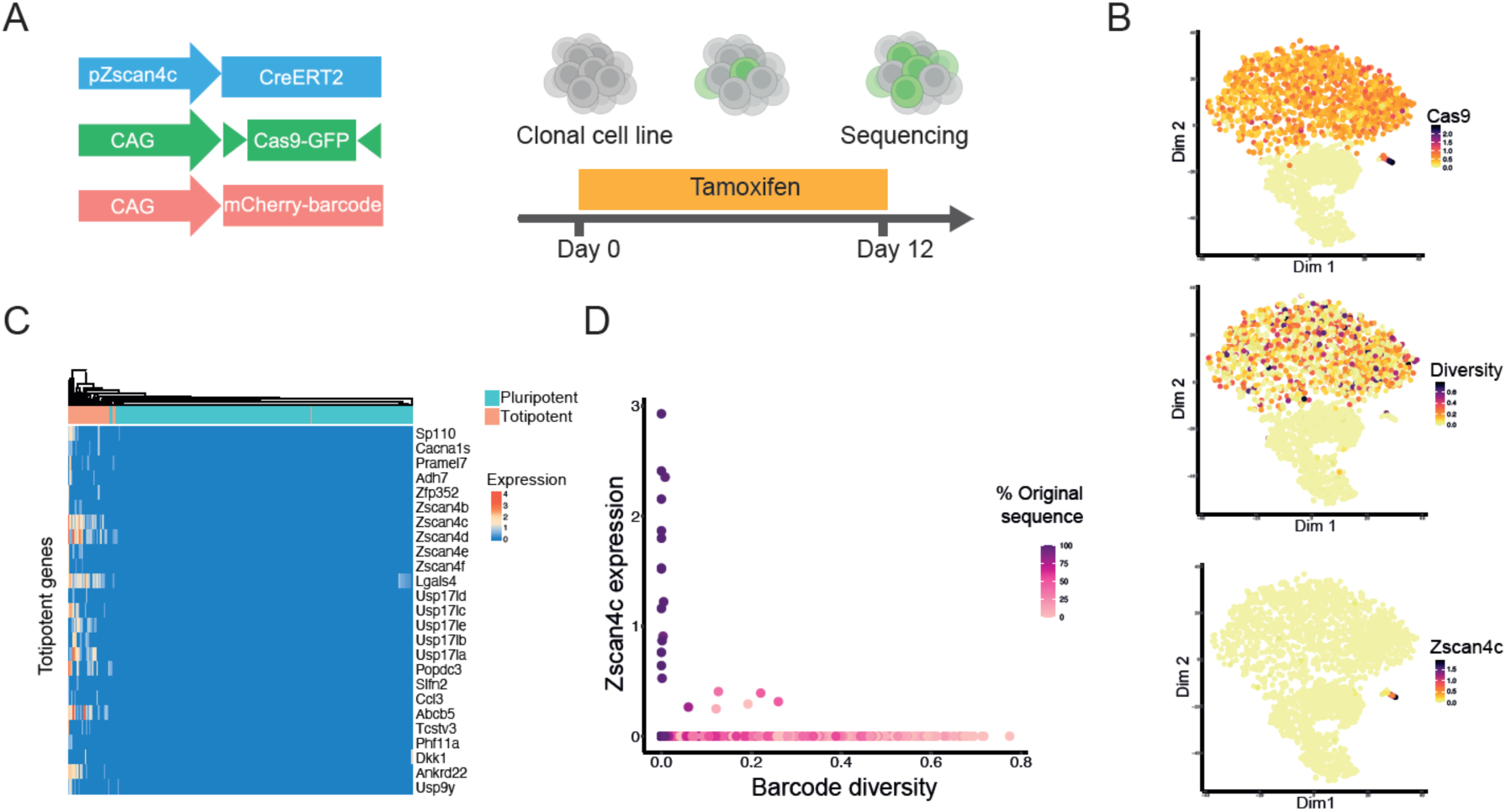
**A.** Schematic representation of the experimental setup for tracking 2C-like state transitions. **B.** t-SNE plot coloured by (top) Cas9 expression, (middle) mean barcode diversity per cell, and (bottom) Zscan4c expression (log2 normalised counts).**C.** Heatmap displaying gene expression (log2 normalised) for top markers for the totipotent cluster. **D.** Scatterplot displaying Zscan4c expression (log2 normalised) (y-axis) against barcode diversity (x-axis) per cell, with colour indicating the percentage of original barcode sequences.

After cell and barcode quality control, we recovered 3,352 GFP-positive cells and 1,270 Cas9-GFP-negative cells (barcode recovery rate = 98%). t-SNE analysis identified three clusters corresponding to Cas9-GFP-negative cells and two clusters representing Cas9-GFP-positive cells. Within the Cas9-GFP-positive population, we observed variable levels of barcode diversity, indicating the activation of barcoding at distinct time points during our time course (**Figure 4B**). t-SNE analysis also revealed a cluster of 24 cells corresponding to 2C-like cells, characterized by expression of totipotency markers, such as Zscan4c (**Figure 4B and 4C).**

Cells in the pluripotent state, characterized by a lack of Zscan4c expression, exhibited high variability in their barcode diversity (**Figure 4B**) (Diversity Brown-Forsythe Test; p = 0.01; Original barcode sequence Brown-Forsythe Test; p = 0.0006). This high variability indicates that the CRISPR barcoding system was activated early upon entry into the 2C-like state, at different time points in each of the cells. As a result, the continuous activity of the barcoding system led to an accumulation of different levels of diversity over time. In contrast, all cells in the 2C-like state (i.e., with Zscan4c expression) displayed low barcode diversity, along with a high proportion of the original barcode sequence (**Figure 4D**) (Diversity Mann-Whitney-Wilcoxon test; p = 0.0007; Original barcode sequence Mann-Whitney-Wilcoxon test; p = 0.0004). This low diversity cannot be attributed to reduced coverage of the barcode cassette in this population (**Supplementary Figure 8**). The absence of 2C-like cells with high barcode diversity indicates that they transitioned into this state only recently. These findings prove the prevailing hypothesis in the field that ESCs undergo continuous dynamic transitions into and out of the 2C-like state, a phenomenon that has been difficult to measure directly until now (*35*, *39*, *41*). Our results also indicate that the 2C-like cells that became pluripotent did not re-enter into the 2C-like state during the experiment. This could be attributed to a combination of factors: probability, as only about 1% of the cells are in the 2C-like state at any given time, and potential inhibitory or promoting signals that might regulate re-entry into this state.

To identify candidate genes that may regulate these transitions, we performed a correlation analysis between the transcriptome profiles and diversity scores in pluripotent cells. Notably, Dppa4, a gene implicated in the 2C-like state emerged as one of the top correlated genes. (**Supplementary Table 3, Supplementary Figure 9**). These findings suggest that the progressive upregulation of Dppa4 in pluripotent cells may facilitate the transition toward the 2C-like state. Indeed, previous studies have demonstrated that Dppa4 depletion inhibits the 2C-like reprogramming in mESCs (*42*, *43*). Other genes correlated with the diversity score potentially involved in the 2C-like transition include the Jumonji Domain Containing 8 (*JMJD8*) or Zinc Finger Protein 226 (*ZNF226)* **(Supplementary Figure 9).**

These results demonstrate the effectiveness of scDynaBar in capturing the dynamic transitions between these states, making it a versatile tool for studying complex cellular dynamics that could not be previously tracked.

### Dynamic barcoding in mouse gastruloids

To investigate the mutation rate across diverse cell types, we applied scDynaBar to mouse gastruloids, which are three-dimensional structures that mimic early developmental processes and comprise a diverse array of cell types (*44*). For this, we used our previously established mESC monoclonal cell line (**Figure 3**). Two days after Cre induction, mESCs were seeded for aggregation and subsequent gastruloid formation (*44*) (**Figure 5A**). Six days after induction, Day 4 gastruloids were dissociated, and the cells were processed for single-cell RNA and barcode sequencing. These gastruloids exhibited normal morphology and low levels of cell death (**Supplementary Figure 10**). After the cell and barcode quality controls, we recovered 1,057 Cas9-GFP-positive cells (i.e., GFP counts >1) and assigned cell types by mapping RNA expression profiles to a reference atlas of mouse embryos, from E6.5 to E8.5 (*45*) (**Figure 5B**). This analysis revealed a bias towards the mesodermal lineage, consistent with our previous studies showing that gastruloids often favour mesodermal or ectodermal cell types (*46*). We then computed the proportion of the original barcode sequence and the barcode diversity score for each cell and found no clear differences between cell populations (**Figure 5C, Supplementary Figure 11A**), indicating that the barcoding rate is similar across diverse cell types (Kruskal-Wallis Test; *p* = 0.3557).

**Figure 5.**
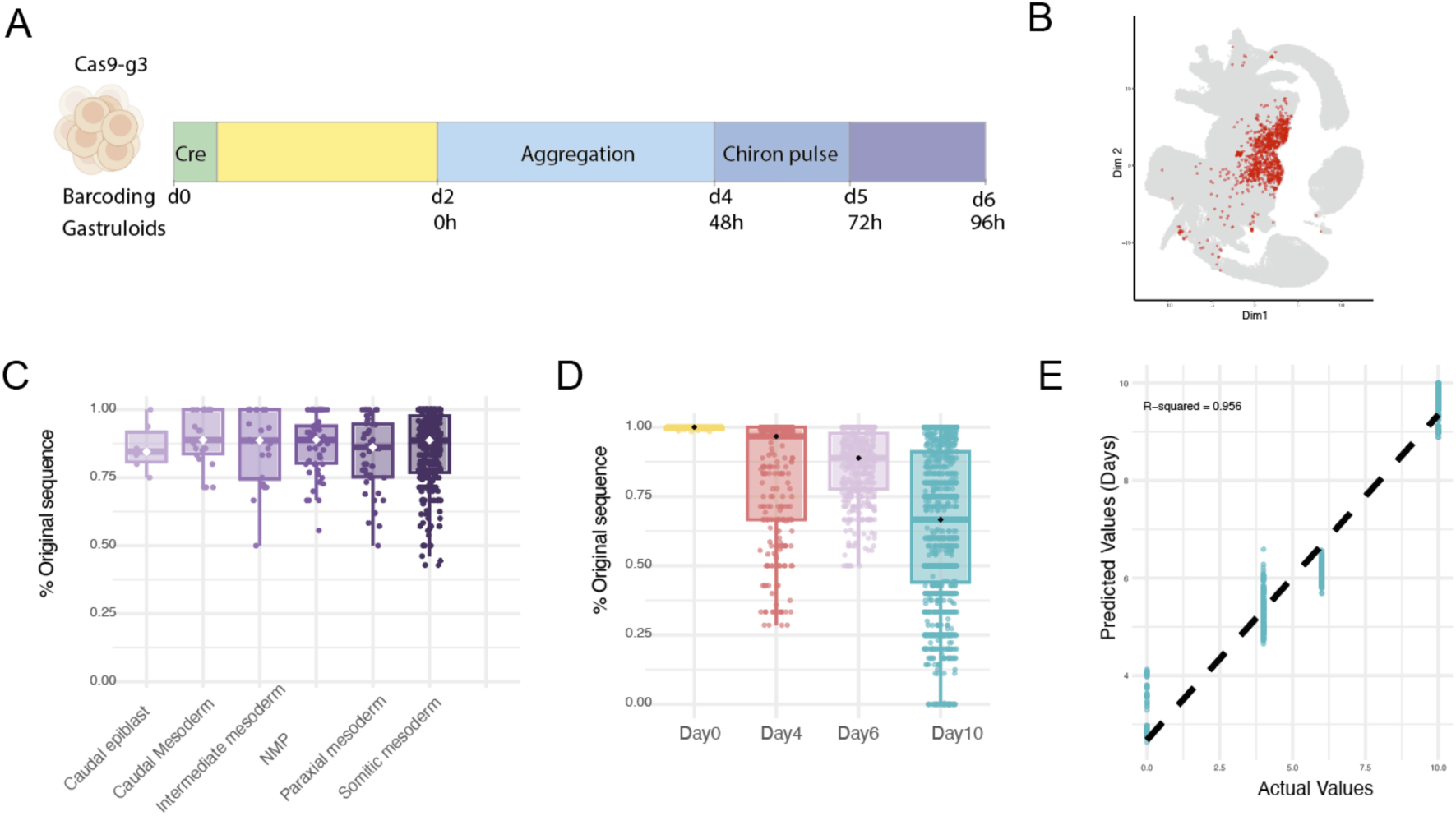
**A**. Schematic of the experimental design for gastruloid formation and barcoding. Cre induction was performed on day 0 (d0). After two days (yellow bar), cells were plated for aggregation (light blue). On day 4 (d4), a Chiron pulse (CHIR99021) was applied to promote gastruloid elongation and cell differentiation (dark blue). Gastruloids were then further cultured (purple bar) and collected on day 6 (d6) for single-cell sequencing **B**. UMAP projection showing developmental stages from a reference atlas of mouse embryos between E6.5 and E8.5, with extra-embryonic cells excluded. Gastruloid cells are highlighted in red. Nearest neighbour analysis was used to assign cell-type labels to the gastruloid cells based on their proximity to the cells in the reference atlas. **C**. Boxplot showing the percentage of the original barcode sequence was calculated for each cell grouped by cell type. **D**. Boxplot showing the distribution of the proportion of original barcodes across different time points in single cells. **E**. Scatter plot showing predicted (y) versus actual (x) time points (days) using a random forest model, with R-squared values of 95% for test data. For all boxplots, the boxes represent the IQR with the horizontal line inside each box indicating the median.

Comparing these data with our previous single cell dataset (**Figure 3**), we observed that gastruloids collected at day 6 exhibited levels of barcoding that ranged between those observed at day 4 and day 10 (**Figure 5D, Supplementary Figure 11B).** Specifically, the median value of the original barcode sequencing was 85% at day 6, compared to 96% at day 4 and 66% at day 10, indicating an accumulative increase of mutations over time (Jonckheere-Terpstra test; *p* = 2 × 10^−04^; 1000 permutations) (**Figure 5D**). We hypothesized that the gradual increase in barcode editing could be used to determine the duration of barcode activity in single cells. To test this hypothesis, we developed a temporal predictor using random forest regression on our single cell datasets (days 0, 4, 6 and 10). The predictor was trained on 80% of the data, incorporating barcode features as variables (mean barcode length, percentage of original sequences, diversity mean and coverage). Using this predictor, we achieved an R^2^ of 98% on the training set, and of 95% on the test data (**Figure 5E**). Achieving this high level of accuracy despite using a limited number of time points suggests that DNA barcodes can effectively be used to predict the duration of barcoding and record the duration of an event in single cells. This highlights the strong potential of dynamic barcoding to function as a molecular clock for time tracking in different cellular contexts.

## DISCUSSION

In this study, we introduce scDynaBar, a novel approach for dynamic cellular barcoding in single cells. We achieved this by engineering self-targeting gRNAs (*31*), enabling the simultaneous capture of both genetic barcodes and the transcriptomes of individual cells. This method requires minimal adjustments to existing commercial single-cell sequencing workflows offering a high level of both versatility and ease of scalability.

We demonstrated that dynamic barcoding leads to the sequential accumulation of genetic diversity over 25 days through continuous targeting, with high reproducibility. This represents a substantial advancement as compared to other single-cell barcoding methods, in which mutations typically occur within the first 48 h (*22*, *26*). The accumulation of mutations over time can be assessed through multiple measures, including i) a reduction in the proportion of original sequences, ii) a decrease in the frequency of sequences with an intact PAM motif and iii) a notable increase in both the diversity of spacer sequences and iv) the variability of their lengths. Interestingly, we observed comparable rates of mutagenesis and barcode diversity using standard Cas9 or the modified cytosine base editor BE3 (*32*), which fuses APOBEC with nCas9. Although potential adverse effects of BE3 were observed during prolonged barcoding, which require further investigation, its efficacy was demonstrated within a 2-week period. This system could be particularly valuable for applications that are incompatible with the generation of double-strand DNA breaks.

As a proof of concept for scDynaBar’s potential applications, we demonstrated its ability to record dynamic cell transitions. By collecting data at a single endpoint and using a CreERT2 protein controlled by the Zscan4c promoter, we documented the transition of mESCs from the pluripotent state to the 2C-like state. In this experimental setup, the diversity of barcodes encodes the timing of the cell transitions to the 2C-like state. Our findings showed that all 2C-like cells had recently transitioned to this state, as evidenced by a lack of mutations in their barcodes. These results underscore the transient nature of the 2C-like state and support the current hypothesis suggesting that the 2C-like state cannot self-perpetuate (*35*, *39*, *41*). Additionally, these findings illustrate the potential of our approach for studying complex cellular dynamics that are currently difficult to track with existing methodologies.

Our observations from a mouse gastruloid model revealed normal phenotypes, indicating the preservation of differentiation potential in cells with active barcoding. Additionally, we observed consistent DNA repair rates across different cell types, underscoring the robustness of scDynaBar and its potential for application in a wide range of biological contexts. Nonetheless, further experiments are required to better understand how factors such as variations in Cas9 levels, DNA repair kinetics, and DNA context influence barcode diversity.

Finally, we believe scDynaBar is highly adaptable and easy to implement, making it suitable for recording a wide variety of temporal events in single cells. For example, by modulating Cas9 activity, scDynaBar could be tailored to track the duration of a specific event, such as the activity of a transcription factor or the response to external stimuli. This flexibility opens new opportunities for capturing dynamic cellular signals and processes at the single-cell level.

## MATERIAL AND METHODS

### Vector construction

The barcoding cassette used in this study were constructed by incorporating gBlock (IDT DNA) synthesized DNA fragments, including CAG promoter, mCherry, and scaffold, into a piggyBac backbone. Different gRNAs (spacers), obtained from Sigma-Aldrich, were cloned into this vector. The sequences of these spacers are provided in the **Supplementary Figure 1**. The Cas9-2A-GFP vector used in this study is based on Cong et al. (2013) (*47*) and was cloned into a piggyBac backbone with a one-way genetic switch (FLEX) system. The BE3 system, kindly provided by Alexis Komor (*32*), was further modified by removing the uracil glycosylase inhibitor (UGI) and then incorporating it into a piggyBac backbone as nCas9-rAPOBEC1-2A-GFP in a one-way genetic switch (FLEX) system.

The plasmid carrying CreERT2 under the control of the Zscan4c promoter (pZscan4-CreERT2 cells), as described in Zalzman et al. (2010) (*40*), was generated by VectorBuilder with a puromycin resistance marker.

### Mouse embryonic stem cell culture

E14 mESCs were cultured under standard serum/LIF conditions in DMEM (Gibco, 11995-040) supplemented with 15% fetal bovine serum, 1 U/ml penicillin and 1 mg/ml streptomycin (Gibco, 15140-122), 0.1 mM nonessential amino acids (Gibco, 11140-050), 4 mM GlutaMAX (Gibco, 35050-061), 50 μM β-mercaptoethanol (Gibco, 31350-010) and 103 U/ml LIF (Stem Cell Institute, Cambridge). Cells were maintained at 37°C in a 5% CO_2_ atmosphere on gelatinized tissue-culture plates. Media was changed daily, and cells were passaged every other day using trypsin-EDTA (Thermo Fisher Scientific, 25200056).

Stable cell lines were generated through transfection using Fugene (Promega, E2311), followed by drug selection and fluorescence-activated cell (FAC) sorting. Clonal cell lines were isolated through additional rounds of subcloning.

To determine the number of copies of the barcode construct integrated in the clonal cell line used in Figure 3 and Figure 5, a genomic qPCR was performed. Genomic DNA was extracted from wild-type (wt) mESCs and from the clonal cell line, using the DNeasy Blood and Tissue Extraction Kit (Qiagen). The qPCR was performed using the Brilliant III Ultra-Fast SYBR® Green QPCR Master Mix in a Roche LightCycler® 480 Instrument II following Agilent Technologies®’ guidelines of one 3-minute cycle at 95°C and 40 cycles of 5 seconds at 95°C and 10 seconds at 60°C with the primers below. Samples were tested in triplicates and a negative control with water was included for each primer combination. Three different DNA amounts were tested: 5 ng, 10 ng and 20 ng. For quantification, the 2^−ΔCt^ method was used, using GAPDH as a reference for the relative quantification of both 2n DNA amplicons (ACTB1, ACTB2 and Intergenic) and the integrated barcoding cassette (Barcode 1 and Barcode 2).

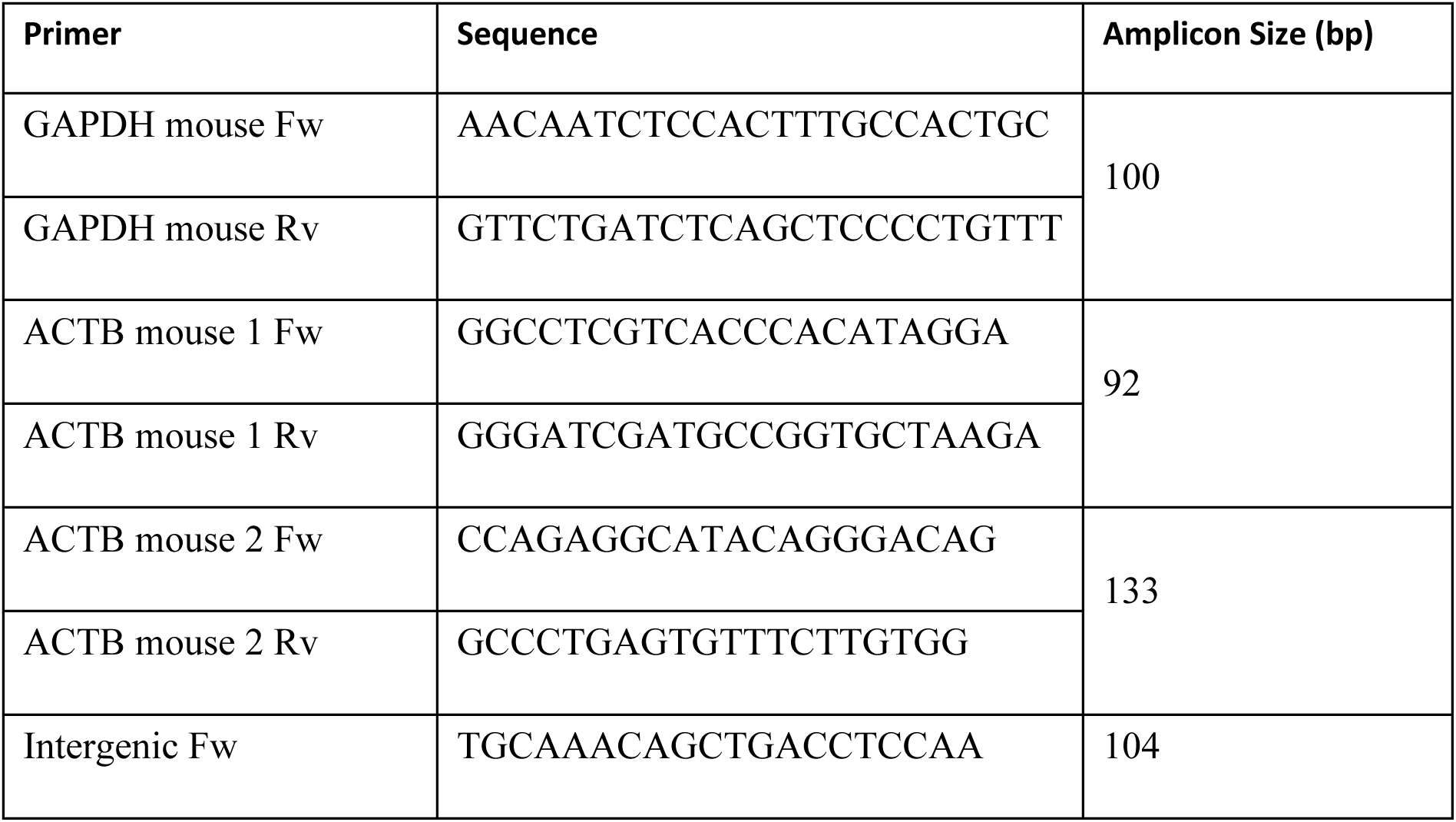
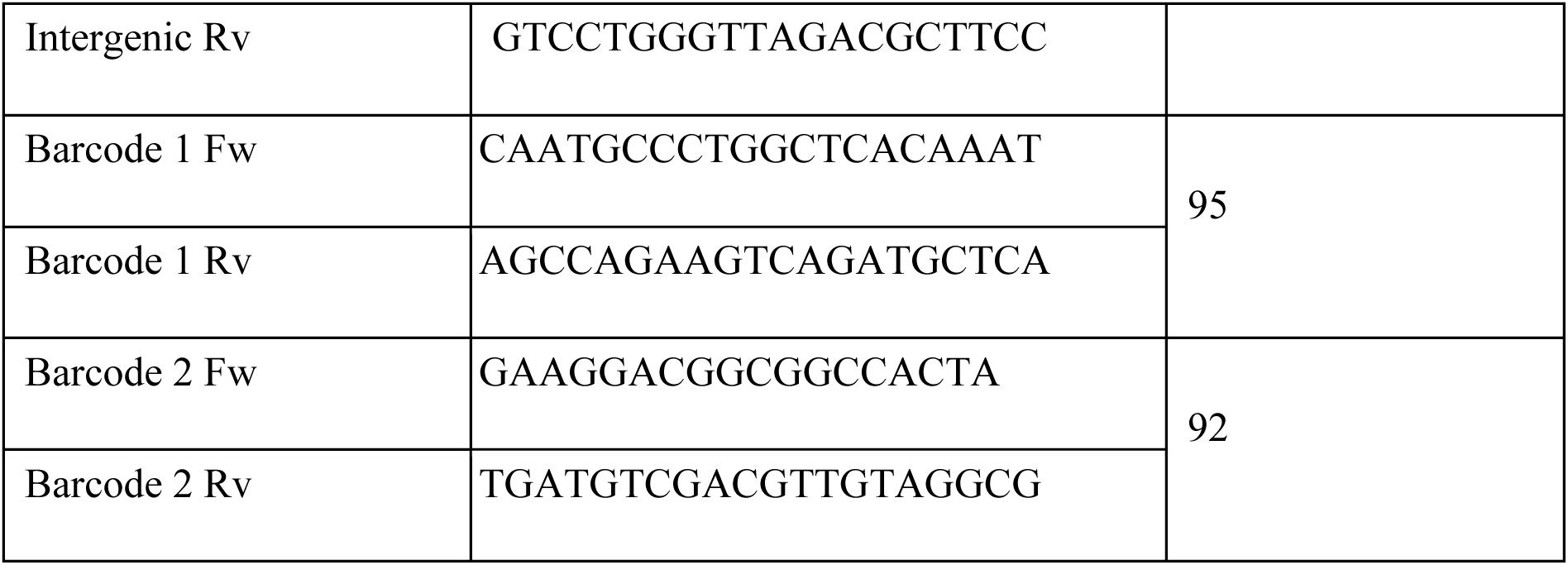

Cas9 induction was carried out using Cre recombinase (Takara, 631449) in media supplemented with 6 µg/mL polybrene (Sigma, H9268), according to the manufacturer’s instructions. pZscan4c-CreERT2 cells were treated with 4-hydroxytamoxifen (1 µM) for 12 days (Sigma, H7904).

### Gastruloid culture

The mESC clonal cell line containing the barcoding system was cultured under standard serum/LIF conditions. Induction was carried out as before using Cre recombinase (Takara, 631449) in media supplemented with 6 µg/mL polybrene (Sigma, H9268), according to the manufacturer’s instructions. After two days, cells were prepared for gastruloid formation using a previously described protocol (*44*). Specifically, mESCs were dissociated into single cells with trypsin-EDTA (Thermo Fisher Scientific, 25300096). Cells were then washed twice with pre-warmed PBS (Thermo Fisher Scientific, 14190144), and the pelleted cells were then resuspended in 5 ml of N2B27 medium of a 1:1 mix of DMEM/F12 (Thermo Fisher Scientific, 11320033) and Neurobasal (Thermo Fisher Scientific, 21103049), supplemented with 0.5× N-2 Supplement (Cell Therapy Systems, A1370701), 0.5× B-27 Supplement (Thermo Fisher Scientific, 17504044), 2 mM GlutaMAX, 10 U/ml penicillin-streptomycin and 0.1 mM 2-mercaptoethanol. Cells were then diluted to a density of 7500 cells/ml in N2B27 medium, and 40 µl of this dilution was added to each well of a U-bottom 96-well suspension culture plate (Greiner Bio-One, 650185) to reach a density of 300 cells per well. The cells were left to aggregate for 48 h. After this period, 150 µl of N2B27 medium containing 3 µM CHIR99021 (Department of Biochemistry, University of Cambridge) was added. Every 24 h, 150 µl of the medium was replaced with fresh N2B27 medium without CHIR99021. On day 4, gastruloids were harvested for sequencing: they were transferred to an Eppendorf tube, rinsed with PBS, and dissociated into single cells using Accutase (StemPro, A1110501). Cells were then washed twice with 5 ml of PBS containing 0.04% BSA (Gibco, 15260037) to remove the Accutase, followed by filtration through a 50 µm strainer (Sysmex, 1050553). The cell count and viability were determined using the Countess II Automated Cell Counter prior to single-cell sequencing.

### Bulk barcode sequencing

Bulk barcode sequencing was performed on a mixed, non-clonal population. After stable integration and induction, cells were sorted at various time points using either the BD Aria III or the BD Influx High-Speed Cell Sorter. About 8K to 12K cells positive for GFP and mCherry were collected for each sample. RNA was then extracted using the RNeasy Micro Kit (Qiagen, 74004). To amplify the gRNA loci, reverse transcription was performed as previously described (*48*), with 1 µl of primer (GTGACTGGAGTTCAGACGTGTGCTCTTCCGATCT-T(30)) and 500 ng of RNA. cDNA was amplified in a 20 µl reaction using KAPA HiFi HotStart ReadyMix (KAPA Biosystems, KK2502) with the following primers:

Forward: 5′-ACACTCTTTCCCTACACGACGCTCTTCCGATCT(N/NN/NNN)TCTTGTGGAAAGGA CGAAACAC-3′

Reverse: 5′-CAAGCAGAAGACGGCATACGAGATXXXXXXGTGACTGGAGTTCAGACGTGTGCT CTTCCGATCT-3′

XXXXXX-index;

Random nucleotides (NNN, NN and N) were added to introduce sequence diversity, enhancing variability over the constant sequence for sequencing purposes. The DNA product was then purified with a 0.8:1 volumetric ratio of AMPure XP Beads (Beckman Coulter, A63881) and the whole volume was resuspended in 20ul of PCR mastermix (KAPA HiFi Readymix) using the primers:

Forward: 5′-AATGATACGGCGACCACCGAGATCTACACTCTTTCCCTACACGACGCTCTTCCGA TCT-3′

Reverse: 5′-CAAGCAGAAGACGGCATACGAGAT-3′

PCR products were purified using a 0.8:1 volumetric ratio of AMPure XP Beads, pooled, and sequenced on a single end Illumina MiSeq run with 58 cycles (58 bp amplicon length) and an 8-cycle index.

### Single-cell sequencing

A single-cell suspension was loaded into the 10X Chromium device, and libraries were prepared using the 10X Single-Cell 3′ Library & Gel Bead Kit v2 (10X Genomics, PN-120237), following the manufacturer’s instructions. The following samples were each loaded into a lane of the 10X Chromium Control chip: day 4, day 10, pZscan4c-CreERT2 cells, gastruloids 1, gastruloids 2, and gastruloids 3.

Amplicon gRNA PCRs (KAPA HiFi Readymix) were performed for each of the 10X cDNA samples using 1 µl of the cDNA and the following primers:

Forward: 5′-ACACTCTTTCCCTACACGACGCTCTTCCGATCT(N/NN/NNN)TCTTGTGGAAAGGA CGAAACAC-3′

Reverse: 5′-AATGATACGGCGACCACCGAGATCTACACTCTTTCCCTACACGACGCTCTTCCGA TCT-3′

As before, random nucleotides (NNN, NN and N) were added to introduce sequence diversity, enhancing variability over the constant sequence for sequencing purposes. PCR products were purified using a 0.8:1 volumetric ratio of AMPure XP Beads, and the whole volume was loaded into a second nested PCR (KAPA HiFi Readymix) with the following primers:

Forward: 5′-CAAGCAGAAGACGGCATACGAGATXXXXXXXXGTGACTGGAGTTCAGACGTGT GCTCTTCCGATCT-3′

Reverse: 5′-AATGATACGGCGACCACCGAGATCTACACTCTTTCCCTACACGACGCTCTTCCGA TCT-3′

XXXXXX-index

After purification using AMPure XP beads, the enriched gRNA libraries were sequenced together with the transcriptome libraries on the Illumina NovaSeq platform.

### Bulk barcode analysis

A total of 160 samples were processed for bulk barcoding (see **Supplementary Table 4**). After sample demultiplexing, reads that did not contain both the flanking regions on the U6 promoter and the spacer regions on either side of the cassette were discarded (“AACAC” and “TTAGAG” respectively). Barcodes with mean Illumina Phred scores below 28 were also removed. Additionally, samples with fewer than 200 reads were excluded. After this quality control, a total of 151 samples were processed for further analysis.

The percentage of original sequences was determined by comparing the spacer regions to the original spacer sequences. To assess diversity, each barcode was aligned to the reference spacer using the *pairwiseAlignment* function from the Biostrings R package (*49*). The alignment was performed using a global-local approach with a substitution matrix that assigned a score of –1 for mismatches and +2 for matches, and a gap opening and extension penalties set to 5. The diversity score for each sample was calculated as the average alignment score across its individual barcodes. Finally, diversity scores were normalized across all samples to ensure comparability. Aligned sequences were further analysed to compute nucleotide frequencies per position.

### Single-cell transcriptome quality control and processing

All 10X scRNA-seq data were processed using the 10X Genomics Cell Ranger pipeline with the mm10 genome build, including the GFP sequence. For quality control, cells were discarded if they had fewer than 3,000 detected genes and/or more than 7.5% of UMI reads originating from mitochondrial genes (**Supplementary Figure 12**). Raw counts for each cell were normalized by dividing the counts by their size factors, and the resulting log-normalized counts were used for further analysis.

Broad clustering analysis and t-distributed stochastic neighbour embedding (t-SNE) plot generation were performed using Seurat. Linear dimensional reduction was executed via principal component analysis (PCA) using the Seurat function RunPCA, with highly variable genes identified by the Seurat function FindVariableGenes as inputs. Fifteen principal components were utilized to generate clusters and t-SNE plots using the Louvain algorithm implemented in the Seurat function FindClusters.

### Gastruloid cell type assignment

Cell types were assigned by aligning RNA expression profiles to a reference atlas as previously described (*46*, *50*). Briefly, count matrices from both datasets were combined and normalized. Highly variable genes were identified and used for PCA. Batch correction was performed to eliminate technical differences between the query and atlas cells. Using the integrated data, a k-nearest neighbours (kNN) graph was constructed. Each query cell’s type was assigned by determining the most common cell type among the 30 nearest neighbours in the atlas, using a Dirichlet distribution for majority voting.

### Single-cell barcode library processing

Fastq single-cell barcode libraries were processed to match cell IDs from the scRNA-seq transcriptomes. Reads lacking the specified flanking regions on the U6 promoter and spacer regions (“AACAC” and “TTAGAG” respectively) or those with an Illumina Phred score of <28, were discarded (**Supplementary Figure 12**). Furthermore, for each UMI with at least three barcode reads, a consensus barcode sequence was determined by selecting the sequence present in over 50% of the retained UMIs. Diversity scores were calculated as in bulk analysis by performing a pairwise alignment of each barcode with the gRNA spacer, using the pairwiseAlignment function from the Biostrings package (*49*). The diversity score for each cell was calculated as the average alignment score across its individual barcodes. Finally, diversity scores were normalized across all cells to ensure comparability.

### Random forest model

The random forest model was employed to evaluate whether barcode editing over time could predict barcode duration in single cells. Single-cell datasets were used from days 0, 4, 6 and 10, whereby the mean barcode length, percentage of original sequences, mean diversity and coverage were incorporated as features. For each dataset, 20% of the data was set aside for testing, while the remaining 80% was used for training. The randomForest function from the randomForest package (version *4.7-1.1*) was used to fit a regression model (*51*), with “day” as the response variable and the selected features as predictors. Model performance was assessed by predicting day values for both the training and test sets, and the R^2^ statistic was calculated to quantify the proportion of variance explained by the model.

## DATA AVAILABILITY

Sequencing data have been deposited in the Gene Expression Omnibus (GEO) database under the accession numbers GSE280613 and GSE280614 (token for revisions are: axavckacjfgzbez and inqlyiuilbgbtqv). The scripts, processed data and metadata are accessible on the GitHub repository: https://github.com/socyol/scDynaBar.git.

